# From Kernel to Kitchen: *Asparagine Synthetase 3* is Involved in Free Asparagine Accumulation in Maize Kernels

**DOI:** 10.64898/2026.06.18.731203

**Authors:** Sarah L. Fitzsimmons, Vivek Shrestha, Abou Yobi, Ruthie Angelovici, Sherry Flint-Garcia

## Abstract

Acrylamide is a probable dietary carcinogen formed through high-temperature cooking processes from one of its precursors, free asparagine (Asn). To reduce acrylamide formation potential in maize-based food products, the genetic basis for the accumulation of free Asn in maize kernels must be elucidated. In this study we integrated complementary quantitative genetic approaches including quantitative trait locus (QTL) mapping of a biparental population and genome-wide association studies (GWAS) in a diverse inbred panel. We ultimately identified one major QTL for free Asn which was not reflected in the GWAS results. Subsequent sequence analysis of the QTL mapping population parents revealed a 921bp deletion in *Asparagine Synthetase 3* (*ASN3*) in the low Asn parent. Validation is underway with rtPCR, genotyping the association panel for the deletion, and evaluation of a CRISPR-Cas9 knockout of *ASN3*. This work provides a foundation for breeding and genome-editing strategies to reduce acrylamide-forming potential in maize-based foods.

## 1. Introduction

Acrylamide is a class 2A, or probable, human carcinogen formed during the Maillard reaction (Mucci and Wilson, 2008) (Figure1C). Under high temperature, low moisture cooking conditions, amino groups rapidly react with reducing sugars to produce desirable Maillard flavor and aroma products. However, other Maillard reaction byproducts react with free asparagine (Asn) to form acrylamide. Since acrylamide’s discovery in 2002, there has been increased vigilance towards reducing its accumulation in thermally processed foods (Halford and Curtis, 2016). This increased scrutiny has prompted the European Union to set standards for food producers (Commission, 2013), and there has been legal action taken against those who do not properly label products containing acrylamide(Sidley, 2025). Maize-based food products such as tortilla chips and breakfast cereals pose concern for dietary acrylamide exposure, especially in children who consume larger quantities of these foods(Mucci and Wilson, 2008).

There have been significant efforts to reduce acrylamide formation potential through metabolic precursors at the plant level. Efforts to reduce sugars have been successful in potatoes (Chawla et al., 2012), but may be less effective in maize, where approximately 70% of the kernel is starch(Watson, 2003, Flint-Garcia et al., 2009), a source of reducing sugars (Žilić et al., 2022). As a result, targeting reducing sugars in maize could significantly compromise kernel size and quality. Targeting free Asn may be deleterious, as it plays a central role in nitrogen storage, transport, and metabolism in plants (Lea et al., 2007). Asparaginases in plants break down free Asn into a usable form of nitrogen to be used for various developmental processes and stress responses (Figure 1B), and because of this, efforts to reduce Asn should be approached with caution (Oliver et al., 2024) to avoid agronomic penalties such as reduced germination (Raffan et al., 2021), yield (Luo et al., 2018), or protein content (Lohaus et al., 1998).

**Fig. 1.**
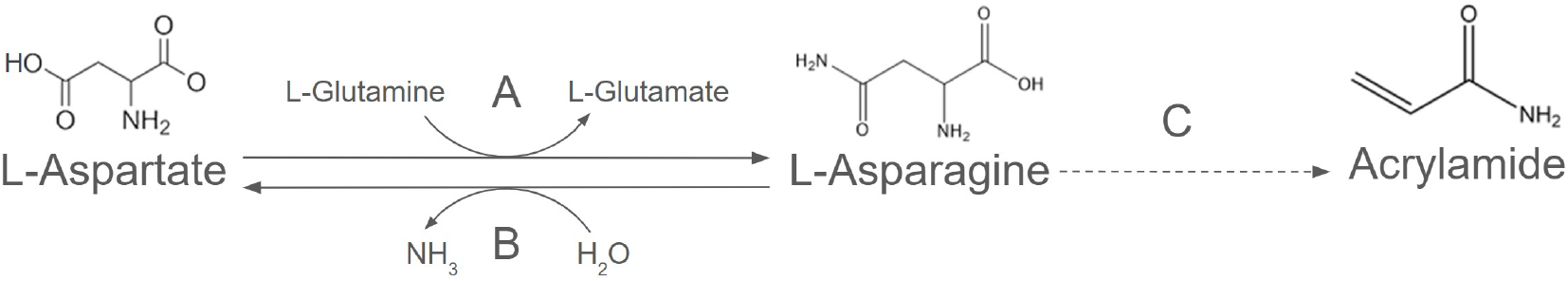
Asparagine (Asn) Metabolism and the Formation of Acrylamide. (A) The metabolism of Asn from aspartate and (B) the degradation of Asn into aspartate and ammonia by asparaginase are among the primary reactions in nitrogen cycling. (C) The formation of acrylamide from Asn is a byproduct of the Maillard Reaction.

In this study, we employed complementary quantitative genetic studies to understand the genetic basis of free Asn accumulation in maize kernels. Quantitative trait locus (QTL) mapping tends to provide robust detection of trait-associated regions(Dhingani et al., 2015), though at lower resolution, while a genome-wide association study (GWAS) offers finer resolution but often has low statistical power(Clauw et al., 2025). Using both methods can provide a more complete understanding of free Asn accumulation. To further refine candidate regions, we constructed whole-genome assemblies for two phenotypically extreme lines. With these combined data, we aimed to unravel the genetic basis of free Asn in maize to provide a foundation for future breeding and biotechnological efforts to reduce acrylamide formation potential in maize-based food products.

## 2. Materials and Methods

### 2.1 Germplasm, Growth Conditions, and Grain Sample Generation

#### Association Mapping Panel

The Goodman-Buckler maize association panel (Flint-Garcia et al., 2005) was grown at the University of Missouri Genetics Farm near Columbia, Missouri in 2017 and 2018, with two replicates each year, in a randomized complete block design. For each plot, 13 kernels were planted in a 10-foot row and self-pollinated to produce grain. At full physiological maturity, the pollinated ears were harvested, husk leaves were removed, and ears subsequently dried at 41 degrees Celsius and 20% relative humidity for 5 days to remove moisture from the grain. Of the 282 lines in this panel, only 279 successfully produced self-pollinated grain. Ears from each plot were shelled and bulked to form four representative composite grain samples for each of the 279 lines in each replicate/year combination. For phenotypic evaluation, 25 random seeds were selected from each bulked sample, ground, and prepared for amino acid extraction and quantification.

#### Quantitative Trait Locus Mapping Population

The bi-parental QTL mapping population was developed by crossing the two inbred lines of the Goodman-Buckler Association Panel(Flint-Garcia et al., 2005) with the lowest and highest free Asn accumulation in the grain, B64 and CML14, respectively, in 2022. The F1 was self-pollinated in winter 2022-2023 to generate the F2 generation, and each F2 plant was self-pollinated in 2023 to produce F2:3 families. The 250 F2:3 families were grown in two replicates of a randomized complete block experiment in 2024 at Genetics Farm near Columbia, Missouri. For each plot, 13 kernels were planted in a 10-foot row and sib-pollinated to produce three ears. Ears were harvested and dried as described above. From each ear, 10 representative kernels were sampled, bulked, ground, and prepared for amino acid extraction and quantification.

### 2.2 Amino Acid Quantification

The free amino acids (FAAs) were extracted and quantified from the grain (Yobi et al., 2020). Briefly, this method utilizes a water-based extraction of amino acids and enables high-throughput detection of all 20 proteinogenic amino acids using liquid chromatography with tandem mass spectrometry (UPLC-MS/MS).

### 2.3 Quantitative Trait Locus Mapping and Analysis

Leaf tissue was collected from F2 plants and lyophilized prior to DNA extraction and genotyping using the mid-density temperate maize single nucleotide polymorphism (SNP) panel by Agriplex Genomics (Cleveland, Ohio, USA). This panel evaluates 2490 SNPs in total and, following filtering of failed, monomorphic, and heterozygous markers, a subset of 611 SNPs was used for the analysis. Markers were re-coded to be compatible with the r/qtl package (Broman et al., 2003) which was used for mapping: AA to correspond when homozygous for the B64 allele, BB if homozygous for CML14, and AB if heterozygous. Best Linear Unbiased Predictions (BLUPs) of 3 Asn traits were calculated:absolute composition (N), relative composition (N.Total), and the ratio of Asn compared to the amino acids of the aspartate family (isoleucine, methionine, Asn, threonine, aspartate, and lysine; N.IMNTDK). These calculations were derived prior to calculating the BLUPs.

Under the advisement of the r/QTL handbook (Broman et al., 2003) and to ensure a high-quality genetic map, markers with more than 10% inconclusive genotyping results were excluded and segregation ratios were checked to identify and remedy potential switched alleles. Linkage groups were formed by calculating the maximum recombination fraction at 0.35 and minimum LOD at 15. Chromosome 5 contained a large low-recombination interval, resulting in two well-supported linkage blocks separated by a wide genetic gap. Pairwise recombination fractions indicated strong internal coherence within each block and minimal recombination between blocks. Several markers in this interval individually inflated genetic map length due to sparse recombination; after removal of the most extreme outliers, remaining gaps were retained as biological features rather than forced into linear order. Ultimately, one linkage group required manual separation and subsequent re-calculation of recombination fraction and LOD to produce an accurate genetic map to reflect the 10 chromosomes of maize. The QTL scans were performed using interval mapping via Haley-Knott regression and significance thresholds were established at a=0.05 using 1000 permutations.

### 2.4 Amino Acid Phenotypic Evaluation and Genome Wide Association Studies

We conducted GWAS for the same 3 Asn traits as QTL mapping: N, N.Total, and N.IMNTDK. For the amino acid GWAS, we used a pipeline based on Holistic Analysis with Pre- and Post-Integration GWAS (HAPPI GWAS) (Slaten et al., 2020). The HAPPI GWAS pipeline removes outliers, optimizes transformation, and calculates BLUPs prior to the association study.

The Goodman-Buckler maize association panel was previously genotyped with both the Illumina MaizeSNP50 Bead-Chip (Cook et al., 2012) and with genotyping-by-sequencing (GBS) (Elshire et al., 2011) methods. We obtained these SNP data from Panzea (www.panzea.org) as the Haplotype Map Version 3 (Bukowski et al., 2018) (HapMap3) dataset, which contained 4,177,796 SNPs. We filtered SNPs by minor allele frequency greater than 0.05 prior to analysis. HAPPI GWAS utilizes GAPIT (Wang and Zhang, 2021) FarmCPU(Liu et al., 2016), MLMM (Segura et al., 2012), and BLINK (Huang et al., 2019) models to conduct univariate GWAS. Following GWAS, we used false discovery rate (FDR) to correct the multiple hypothesis testing problem at 5%. Using the Maize AGP_V4 genome sequence, we identified candidate genes within 100kb on either side of all SNPs that passed FDR correction.

In light of the QTL mapping results, we also performed GWAS using custom VCF (genotypic data) files containing SNPs specific to *ASN3*. We utilized SNP data made available by the SNPVersity tool (Grzybowski et al., 2023) on MaizeGDB (Woodhouse et al., 2021) and produced a custom dataset containing SNPs on chromosome 1 between base pairs 44,907,435-44,914,218 using unimputed short-read sequence data (Bukowski et al., 2018). Following GWAS, we integrated the P-values from the original and custom analyses and re-calculated the false discovery rate correction.

### 2.5 Genome Construction and Analysis

Seed of the QTL parental lines B64 and CML14 was sent to Cornell University where they were germinated, and DNA was extracted from leaf tissue. HiFi reads were produced by PacBio (Menlo Park, California, USA) and used for depth coverage analysis and genome construction. Samtools (Li et al., 2009) was used to align, sort, and index BAM files into compatible formats for visualization in IGV (Thorvaldsdottir et al., 2013). Genome construction was carried out by using the HiFiasm package (Cheng et al., 2021).

### 2.6 Multiple Sequence Alignment

Genomic DNA, coding DNA, and protein sequences for the four maize *ASN*s in the B73 background were sourced from MaizeGDB. Multiple sequence alignment (MSA) was performed using MAFFT v7.525 (Nakamura et al., 2018) and percent identity was calculated using EMBOSS distmat with nucleotide-based distance estimation for genomic and coding DNA sequences and pairwise percent identity was calculated from the protein sequence using EMBOSS distmat.

## 3. Results

### 3.1 Natural Variation in free Asn

The Goodman-Buckler Association Panel exhibited extensive natural variation in free Asn content at grain maturity, with a nearly 20-fold difference between the lowest line (B64; average = 0.172 nMol/µg Asn) and one of the highest lines, CML14 (average = 12.419 nMol/µg Asn; Figure 2A)). This result suggested that not only is it biologically possible for there to be low free Asn corn, but there are genetic controls governing its accumulation in kernels.

**Fig. 2.**
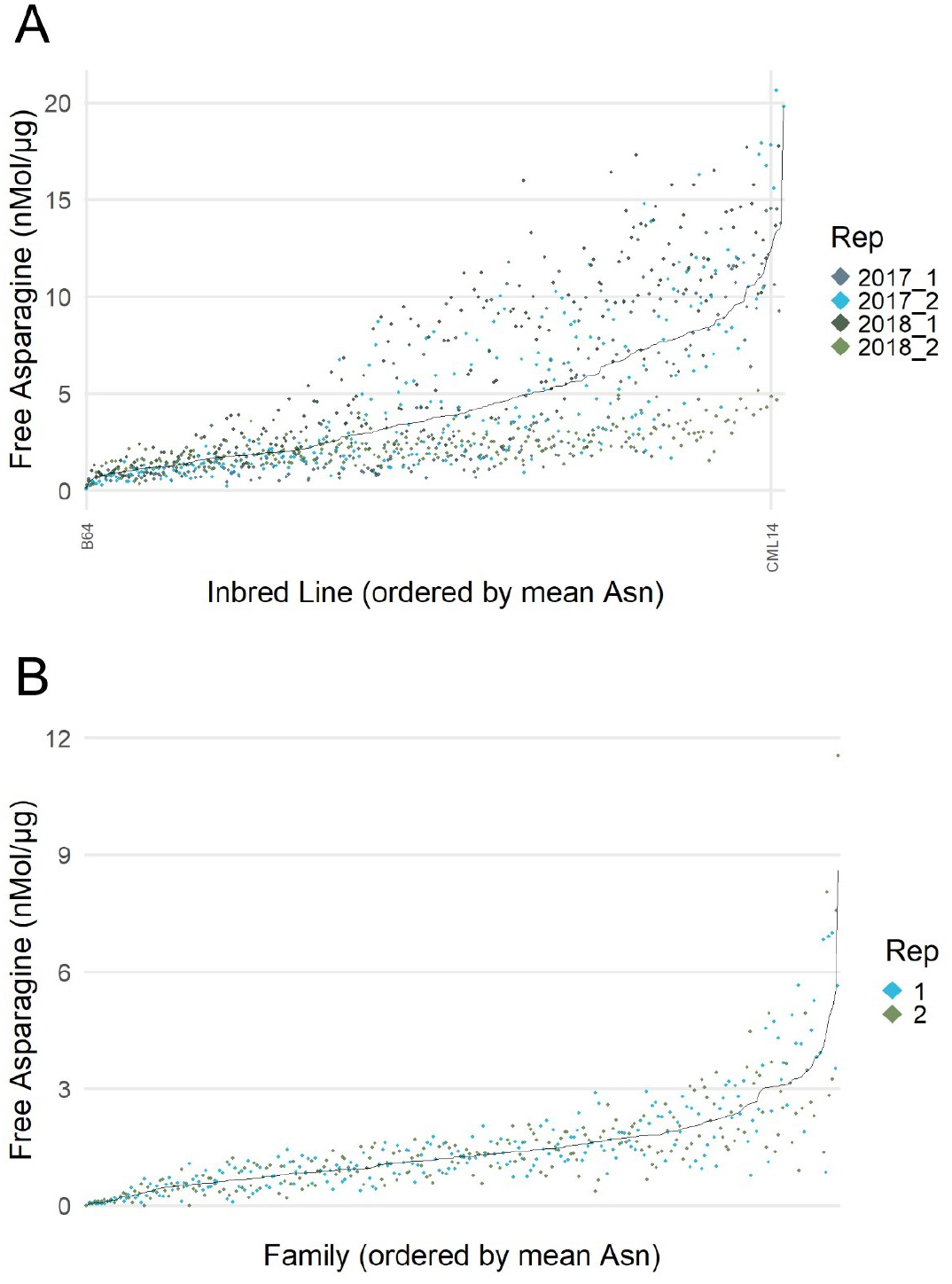
Distribution of Kernel Free Asn in the Two Mapping Populations. Both the (A) Goodman-Buckler association panel and (B) QTL mapping populations show natural variation. Select inbred lines are labeled including the lowest inbred (B64) and one of the highest (CML14) in the association panel population.

### 3.2 Quantitative Trait Locus Analysis

We performed QTL mapping in an F2:3 population derived from B64 × CML14 to identify loci associated with free Asn content in maize kernels. Best Linear Unbiased Predictions of the 3 Asn traits across the two replicates were calculated from raw data (Figure 2B) and used for the interval mapping QTL scan using Haley-Knott regression. The QTL scans were nearly identical for the three Asn composition traits: absolute Asn (total Asn) and Asn relative to the sum of all FAAs (N.Total), and Asn relative to the aspartate family (N.IMNTDK) (Supplemental Figures S1-S3). All future references to QTL mapping results will refer to absolute Asn. The genetic map consisted of 611 SNP markers (Supplemental Figure S4) distributed across 10 linkage groups, and the significance threshold for QTL mapping was determined to be LOD = 3.9 after 1,000 permutations (α = 0.05).

A single, major QTL was identified on chromosome 1 (Figure 3) with a peak LOD score of 18.9. This QTL spanned over 28 markers, equating to base pair positions 22,816,035 through 90,306,830. This QTL explains 29.8% of the variance for this trait and the allele that increased free Asn originated from the high Asn parent, CML14. The LOD support interval (1.5 LOD drop) spans base pairs 39,838,119 through 55,146,349, which aligns with the location of *Asparagine Synthetase 3* (*ASN3*) which is located at chr1:44,908,935-44,912,718. This result aligns with what is documented in other crop species as *ASN*s have been shown to be the primary drivers of free Asn accumulation in plant tissues.

**Fig. 3.**
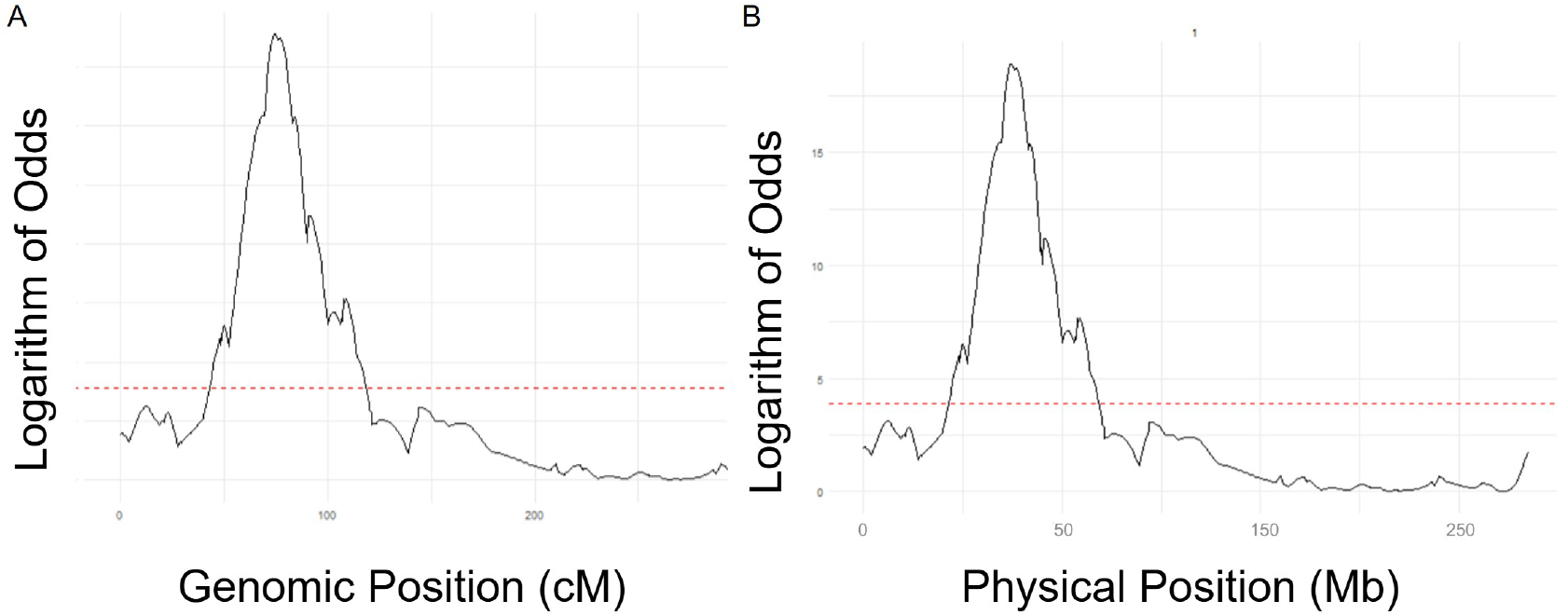
Major QTL for Free Asn. Identified from biparental F2:F3 mapping population on chromosome 1, with red dashed line representing threshold of significance. (A) Genomic position. (B) Physical position.

### 3.3 Genome Wide Association Studies

The FAA content and composition of dry, mature maize kernels were measured from 279 lines belonging to the Goodman-Buckler Association Panel, which was grown in Columbia, Missouri in 2017 and 2018, with two replicates each year. A water-based extraction method was used to extract the 20 proteinogenic amino acids which were subsequently quantified using LCMS/MS detection. These data were used to perform the genome wide association study for the three Asn composition traits.

To identify genetic loci associated with free Asn accumulation in maize kernels, we first conducted genome-wide association studies (GWAS) across the three traits and three models. After applying false discovery rate correction at 5%, 218 unique, significant SNPs were identified across all models and traits (Supplemental Figures S5-S13). As expected, FarmCPU yielded the most results with 165 unique, significant SNPs, followed by BLINK with 45 unique SNPs, and MLMM with only 1 significant SNP. The overlap between models was modest, with 17 SNPs appearing for more than one model (Figure 4A), while only one SNP (3-212191792) was identified across all three models. Most SNPs were unique to the trait they were associated with, with only six being shared between N.Total and N.IMNTDK, and one between N.IMNTDK and N (Figure 4B).

**Fig. 4.**
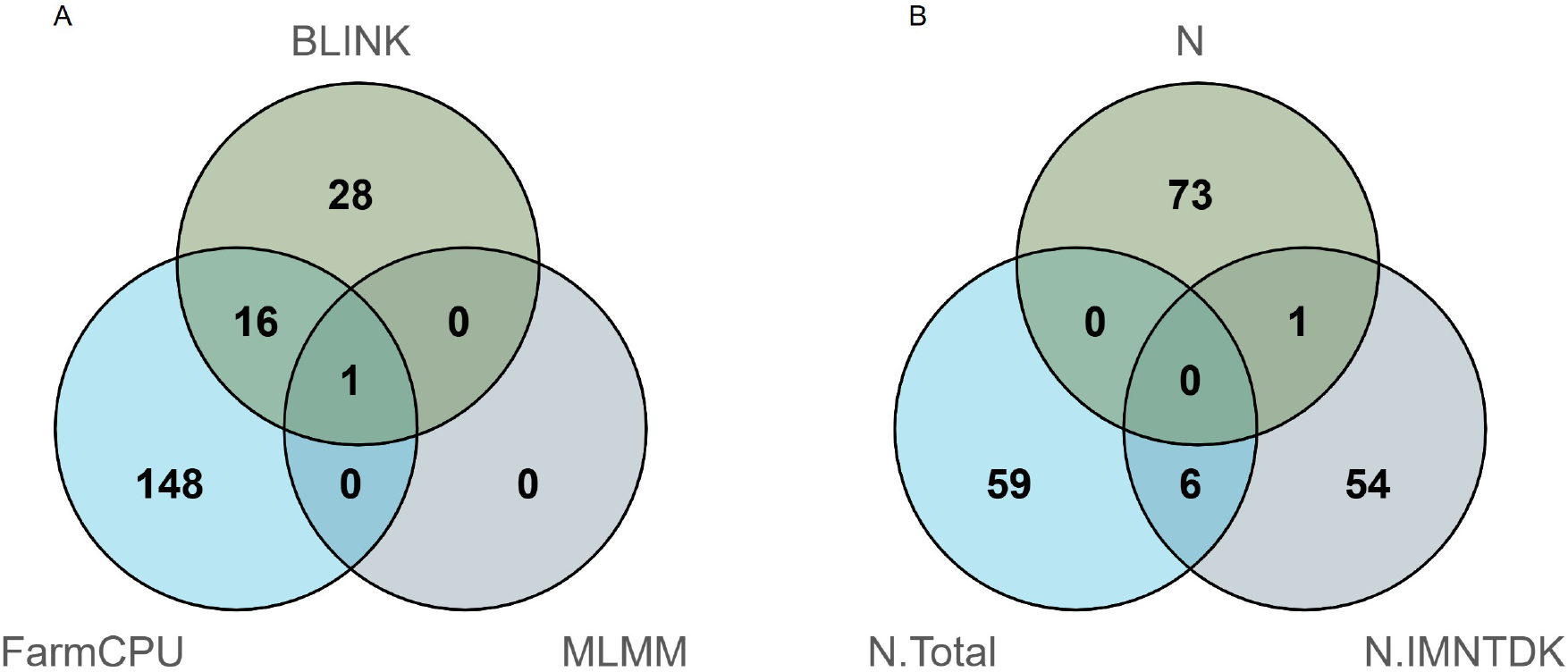
Results of Genome Wide Association Studies. (A) Overlap of significant SNPs identified by model. (B) Overlap of significant SNPs identified by trait.

BLINK, or Bayesian-information and Linkage-disequilibrium Iteratively Nested Keyway, leverages linkage disequilibrium (LD) information to select non-redundant markers when performing stepwise regression (Huang et al., 2019). There was no overlap between the three traits analyzed using BLINK. The 45 unique, significant SNPs identified by BLINK correlated to 1,311 candidate genes within the 100kb window on either side of each SNP. Of the candidate genes, 929 had annotations in MaizeMine, a tool in MaizeGDB. Neither *ASN3* nor any of the other *Asparagine Synthetase* (*ASN*) genes were identified as candidate genes.

FarmCPU, which stands for Fixed and Random Model Circulating Probability Unification, circulates between two models that account for both fixed and random effects (Liu et al., 2016). It produced the most results of this analysis, with a total of 165 unique, significant SNPs which equate to 4,859 candidate genes. Again, *ASN3* was not among the results. However, there were interesting findings with biological significance such as Amino Acid Permease 6 which has been shown to influence free Asn transport into kernels (Wang et al., 2022). FarmCPU also identified many more ribosomal proteins than BLINK.

The final model used, Multi-Locus Mixed Model or MLMM, is the most statistically rigorous of the three and functions by adding multiple SNPs as covariates through stepwise selection (Segura et al., 2012). This model produced a single significant SNP, which correlated to 31 candidate genes. This result was the only SNP to appear for all three models, underscoring its role in free Asn accumulation.

Despite this fact, no annotation of these 31 candidate genes indicated a clear biological role.

Ultimately, a large number of significant SNPs were detected across models, equating to 5,482 candidate genes, which is approximately 10% of all annotated maize genes. Despite these abundant results, no candidates corresponded to *ASN3* nor any other *Asparagine Synthetase*. This absence was unexpected based on biological priors (biochemical pathways and results from other species) and our QTL evidence (Figure 3). Closer inspection of the HapMap3 dataset used for the initial GWAS revealed that there were no SNPs in *ASN3*; therefore, associations at this locus could not have been detected regardless of effect size. This limitation motivated the subsequent efforts to obtain SNPs within the ASN3 region.

The custom VCF dataset contained un-imputed genotypic data with 517 SNPs from chromosome 1, between base pairs 44,907,435 and 44,914,218, corresponding to *ASN3*. GWAS results from this dataset across three models were integrated into the HapMap3-based GWAS results. Following re-calculation of FDR, one SNP from this *ASN3*-enhanced dataset was significant. This intergenic SNP (Chr1:44,907,882), identified by BLINK for the trait N.IMNTDK, provides evidence that *ASN3* contributes to variation in this panel.

### 3.4 Depth Coverage Analysis

To assess structural variation between the QTL parental lines, we performed a read-depth analysis across the four annotated *Asparagine Synthetase* (*ASN*) genes in maize. Coverage was generally uniform and complete for *ASN1* (chr9:34,625,482-34,634,881), *ASN2* (chr3:235,198,390-235,201,558), and *ASN4* (chr9:143,805,569-143,809,317) in both CML14 and B64, indicating no major structural differences at these loci. However, *ASN3* (chr1:44,908,935-44,912,718) exhibited a 921bp deletion in B64 (Figure 5) corresponding to 3 exons that are present in all three transcripts of *ASN3* in the B73 genome, suggesting a potential functional impact on free Asn metabolism.

**Fig. 5.**
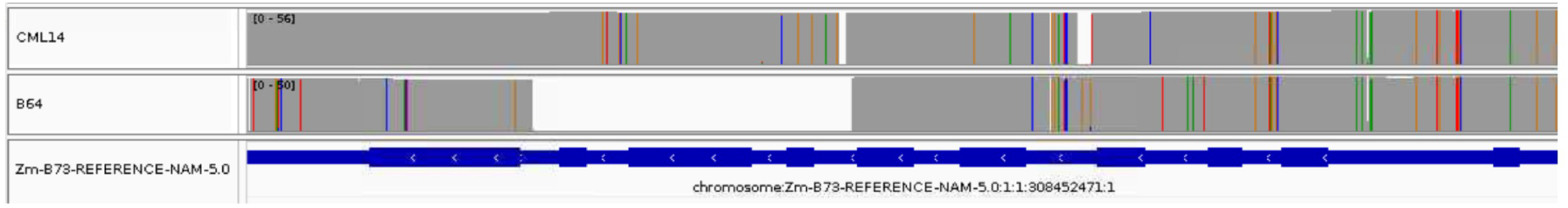
Read Coverage of *ASN3* Showing 921bp Deletion in B64. Colored lines indicate mismatched base pairs where red = Thymine, green = Adenine, blue = Cytosine, and orange = Guanine.

### 3.5 Multiple Sequence Alignment

Multiple sequence alignments were performed for genomic DNA, coding sequences (CDS), and protein sequences of the four *ASN*s in B73. In genomic and CDS alignments, *ASN3* and *ASN4* exhibited negligible pairwise divergence, whereas comparisons between *ASN1* or *ASN2* and *ASN3*/*ASN4* showed markedly greater divergence. Protein sequence alignment similarly revealed high conservation between *ASN3* and *ASN4* (>95% identity), in contrast to lower similarity between these genes and other *ASN* paralogs.

## 4. Discussion

The presence of multiple copies of the *Asparagine Synthetase* (*ASN*) gene is often associated with tissue-specific roles. For example, in potatoes, *ASN1* primarily regulates free Asn accumulation in leaves, whereas *ASN2* controls levels in tubers (Chawla et al., 2012). Similarly, in wheat, *ASN2* shows the greatest specificity to grain (Raffan et al., 2021). In maize, there are four copies of *ASN*. Overexpression of *ASN2* and *ASN3* results in increased free asparagine in kernels (Crowley et al., 2025). Although natural expression of *ASN* genes in B73 kernels is low, *ASN3* exhibits the highest expression among them (Oliver et al., 2024), making it the most plausible candidate of the four in maize for controlling free Asn accumulation. Therefore, our a priori expectation was that we would identify *ASN3* as one of the genes underlying free asparagine accumulation in our study.

Our hypothesis was supported by the QTL mapping results. Though we expected to identify multiple QTLs, which is typical for quantitative traits, we identified a single, major QTL on the short arm of chromosome 1 that controls approximately 30% of the variance in free Asn in maize kernels. This QTL peak located at 44,793,022 bp has a confidence interval of base pairs 39,838,119 through 55,146,349. This confidence interval encompasses *ASN3* (44,908,935–44,912,718 bp), strongly suggesting *ASN3* as the causal gene underlying this locus.

The two parents of the QTL mapping population belong to the Goodman-Buckler Association panel, which is composed of 282 diverse inbreds from across the world. Though QTL mapping is a high-power tool used to identify loci associated with traits, it lacks the resolution that can often be accomplished in GWAS. When performing GWAS using this panel, we expected to find *ASN3* among the results since the two parental lines of the QTL population are members of the association panel. However, this was not the case. This discrepancy between the strong QTL signal for *ASN3* in the biparental population and its absence in the GWAS results was unexpected and prompted further investigation.

Inspection of the HapMap3 dataset 23 revealed that there were no SNPs within *ASN3* nor any of the other *Asparagine Synthetase* genes. Though there are ample SNPs within the 100kb window on either side of *ASN3* (276 and 84 SNPs upstream and downstream, respectively; Figure 6A), none were significant after FDR correction. Linkage disequilibrium on chromosome 1, where *ASN3* is located, decays rapidly (Figure 6B). At the distance between the closest upstream SNP and the start of *ASN3* (approximately 2 kb), LD is moderate (R^2^ = 0.537), and similarly for the nearest downstream SNP to *ASN3*’s end (approximately 8 kb), R^2^ = 0.384 (Figure 6B). These values indicate that nearby SNPs were only weakly correlated with *ASN3*, which sufficiently explains why any potential phenotypic association with Asn was not tagged by the flanking SNPs and absent among the GWAS results.

**Fig. 6.**
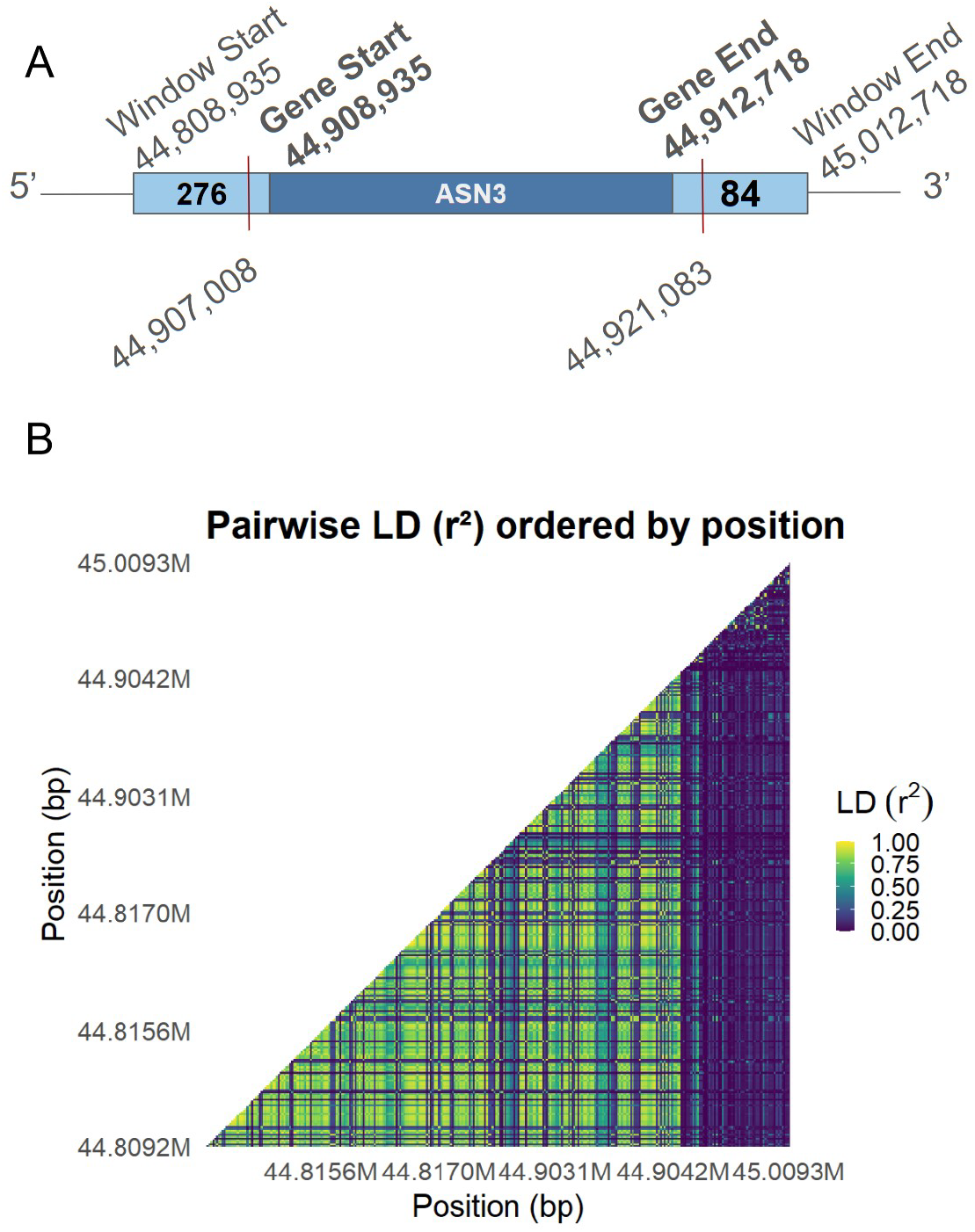
SNP Density and marker coverage that potentially explains the absence of *ASN3* in GWAS results. (A) There 276 SNPs upstream of *ASN3* with the closest being approximately 2kb from the start and 84 downstream of the end, with the nearest SNPs (located at 44,907,008 and 44,921,083 bp) being approximately 8kb away. (B) Linkage disequilibrium decay in 200kb search window surrounding *ASN3*.

To further investigate whether variation within *ASN3* could be detected in GWAS, we incorporated additional SNP data from SNPVersity2.1 and reran the GWAS models. Of the 517 un-imputed SNPs, only one SNP located in the intergenic region was significant after FDR correction.

To investigate potential structural variation as an underlying cause for the discrepancy between QTL mapping and GWAS results surrounding *ASN3*, the QTL mapping population parents were sequenced using the PacBio system. We examined HiFi reads using the Integrative Genomics Viewer (IGV) to assess depth and coverage. This analysis revealed a 921 bp deletion within *ASN3* in B64, spanning three exons that are essential for all three transcripts that are annotated in the B73 genome. Such a deletion likely explains both the pronounced divergence in free Asn content between parental lines and the absence of *ASN3* in the GWAS results. Interestingly, this short-read sequenced dataset contained SNPs within a genomic region corresponding to a deletion identified by long-read sequencing of B64. Given that *ASN3* contains a true deletion in B64, it is likely that the short-read signal mapping to that interval must originate from the highly similar paralog *ASN4*.

These findings provide a foundation for developing maize varieties with reduced acrylamide-forming potential, enabling the production of safer, low-acrylamide maize-based food products. Development of maize germplasm with low free Asn may be achieved by employing genome-editing approaches targeting *ASN3* in commercial lines or by employing marker-assisted selection for the 921 bp deletion allele in breeding programs. A deeper understanding of *ASN3* biology is still needed, and forthcoming gene-editing experiments and RNA-based studies to assess *ASN3* expression and transcript integrity will help clarify the regulatory and functional roles of *ASN3* in kernel Asn accumulation.

## Acknowledgements

S.F. was supported by the Marcus S. Zuber Assistantship and the Oak Ridge Institute for Science and Education CERCA Project. S.F.G. was supported by USDA-ARS. The authors would also like to thank Drs. Daniel Kick and Ha Ngoc Duong for guidance in conducting the GWAS and the members of the Flint-Garcia Lab for their contributions in pollination and harvest of the QTL mapping population. Mention of trade names or commercial products in this publication is solely for the purpose of providing specific information and does not imply recommendation or endorsement by the U.S. Department of Agriculture. USDA is an equal opportunity provider and employer.

## Supplemental Figures

**Fig. S1.**
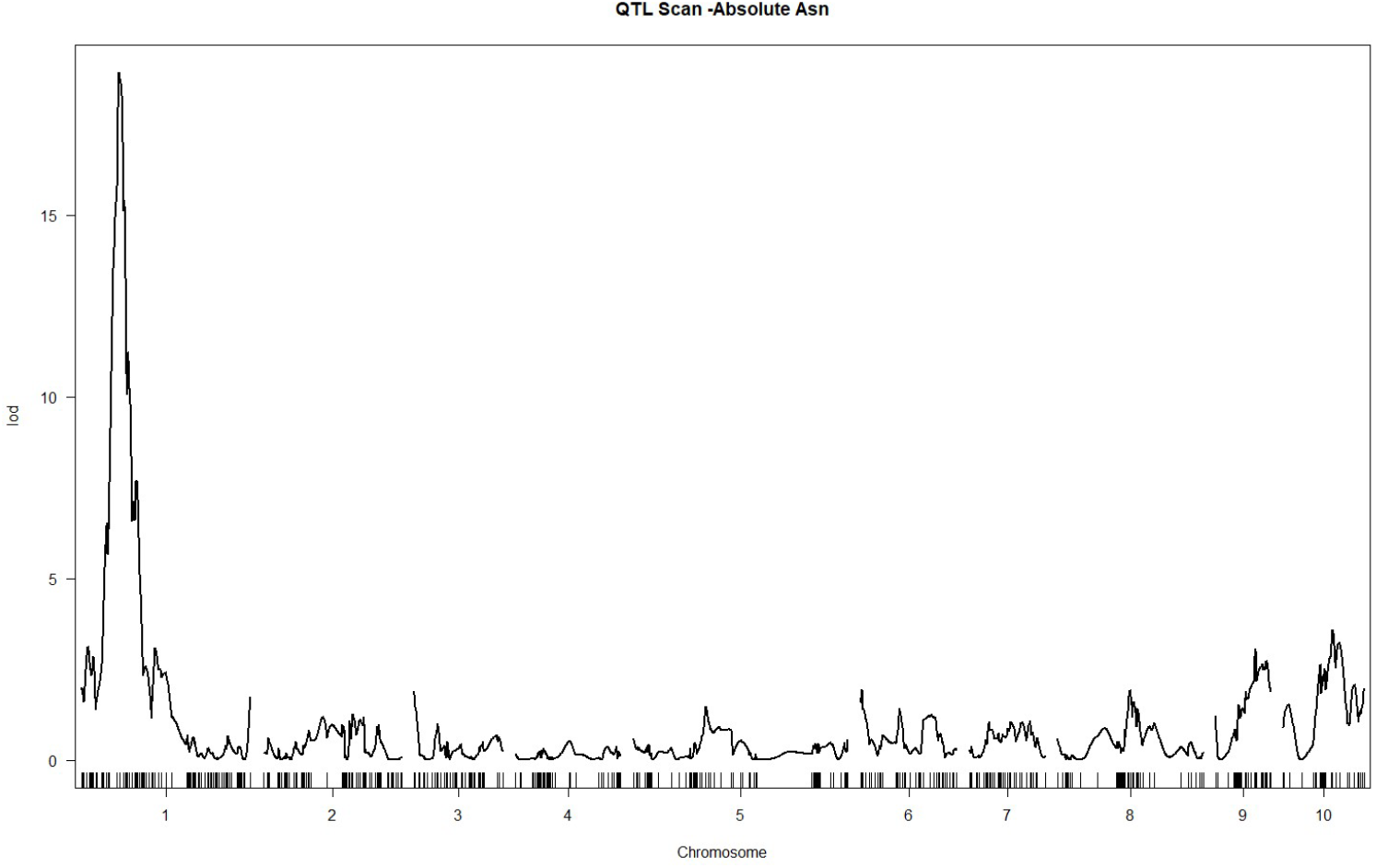
QTL Scan for Absolute Asn

**Fig. S2.**
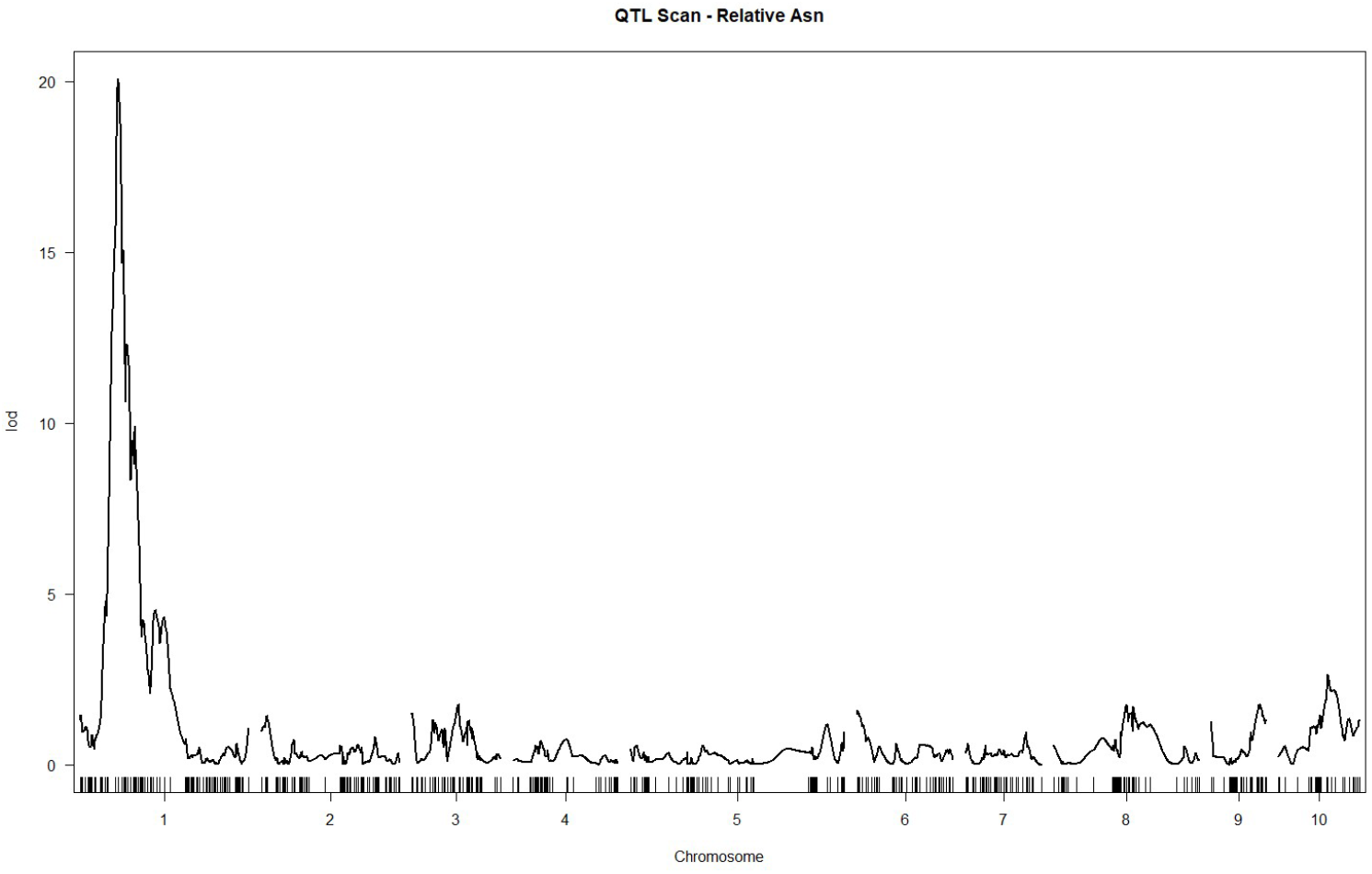
QTL Scan for Relative Asn

**Fig. S3.**
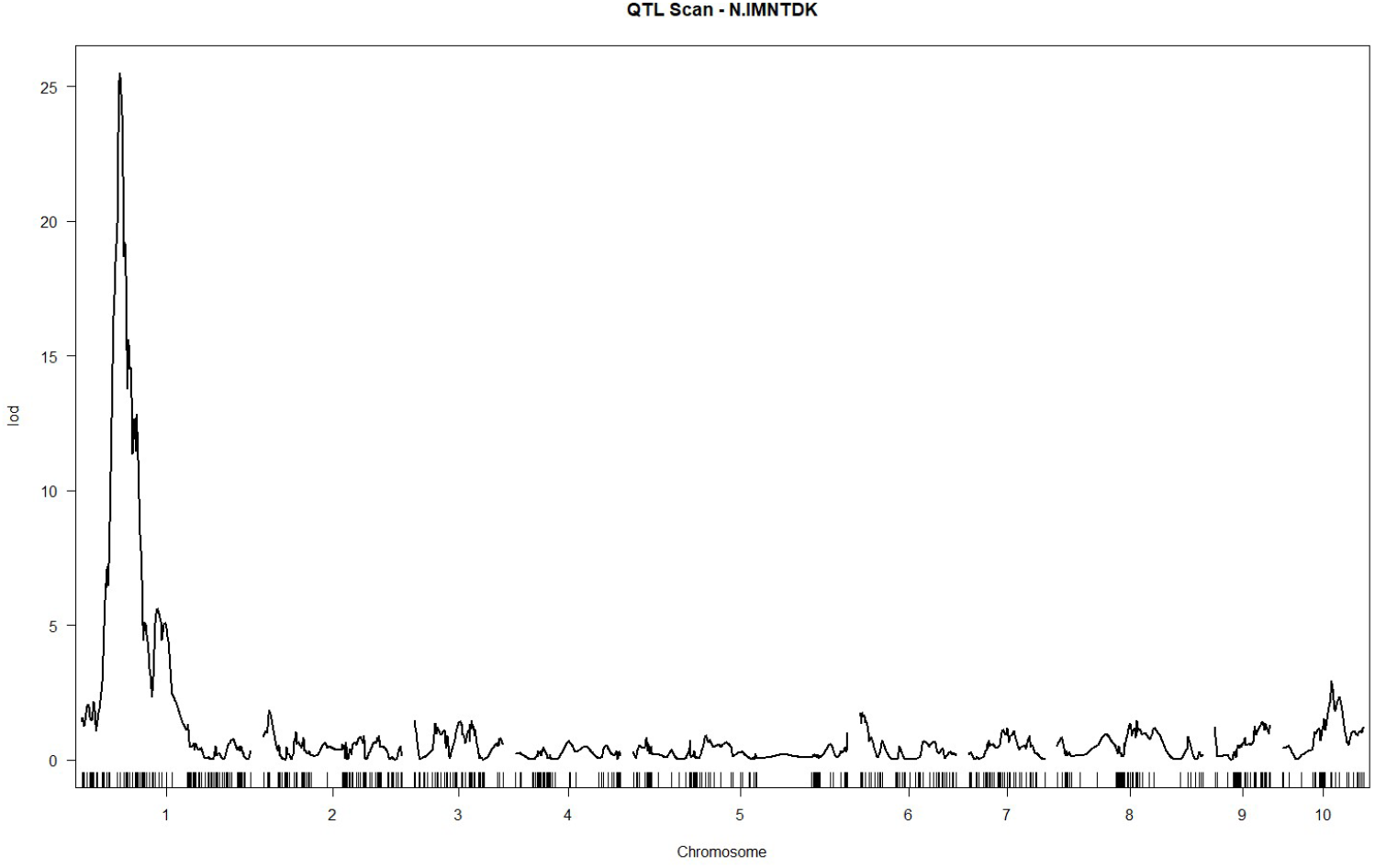
QTL Scan for Family Asn

**Fig. S4.**
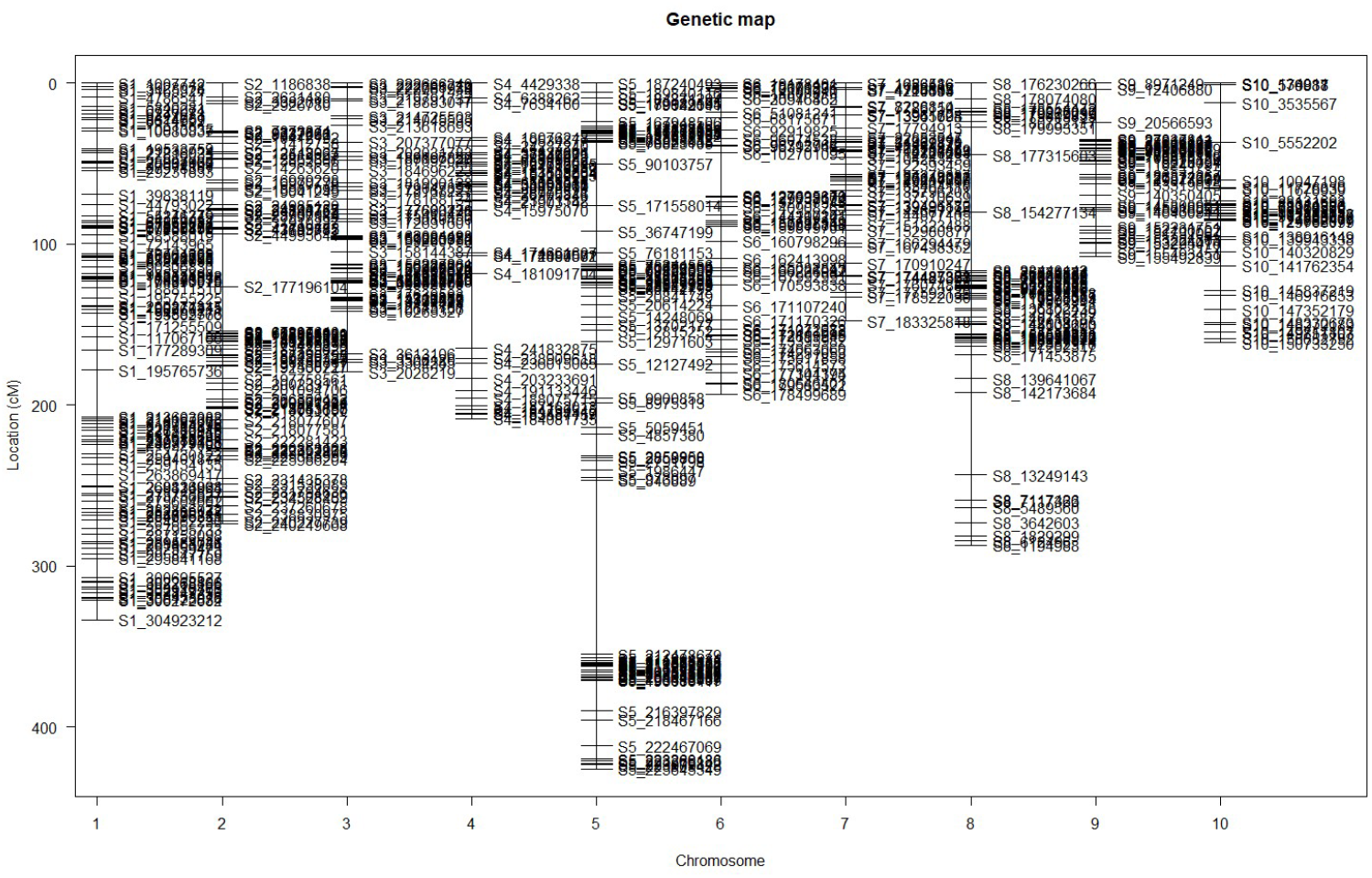
Genetic Map

**Fig. S5.**
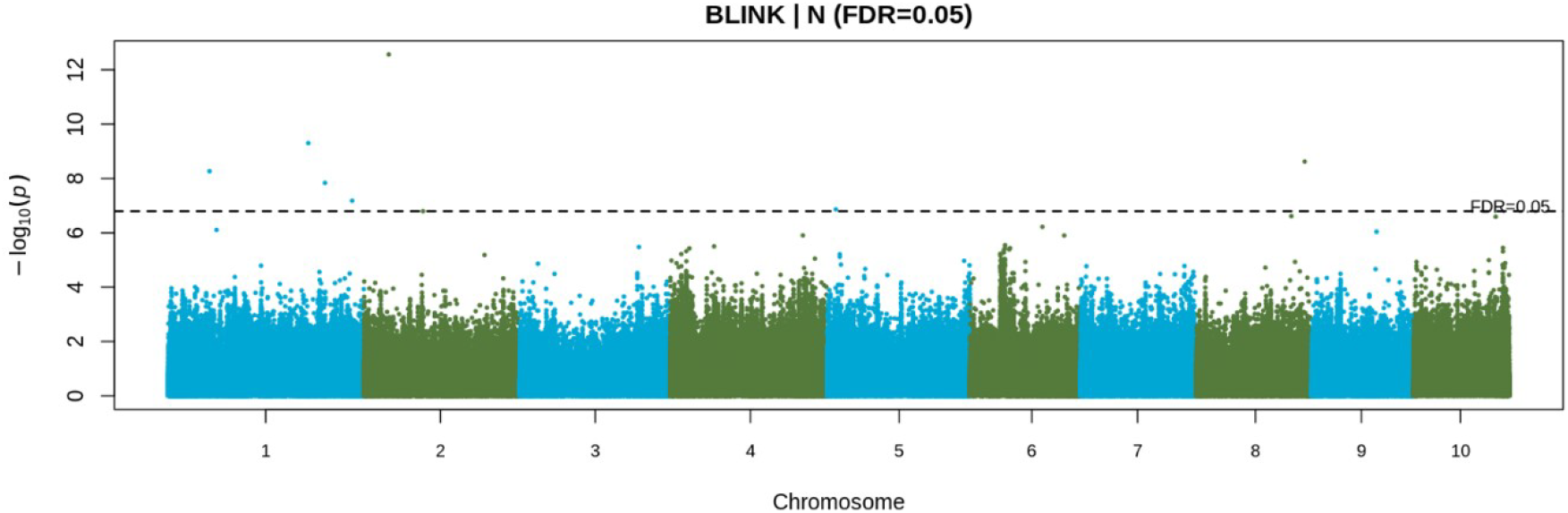
Manhattan plot of BLINK results for absolute free Asn. Points above dashed line represents results that passed false discovery rate correction of 5%.

**Fig. S6.**
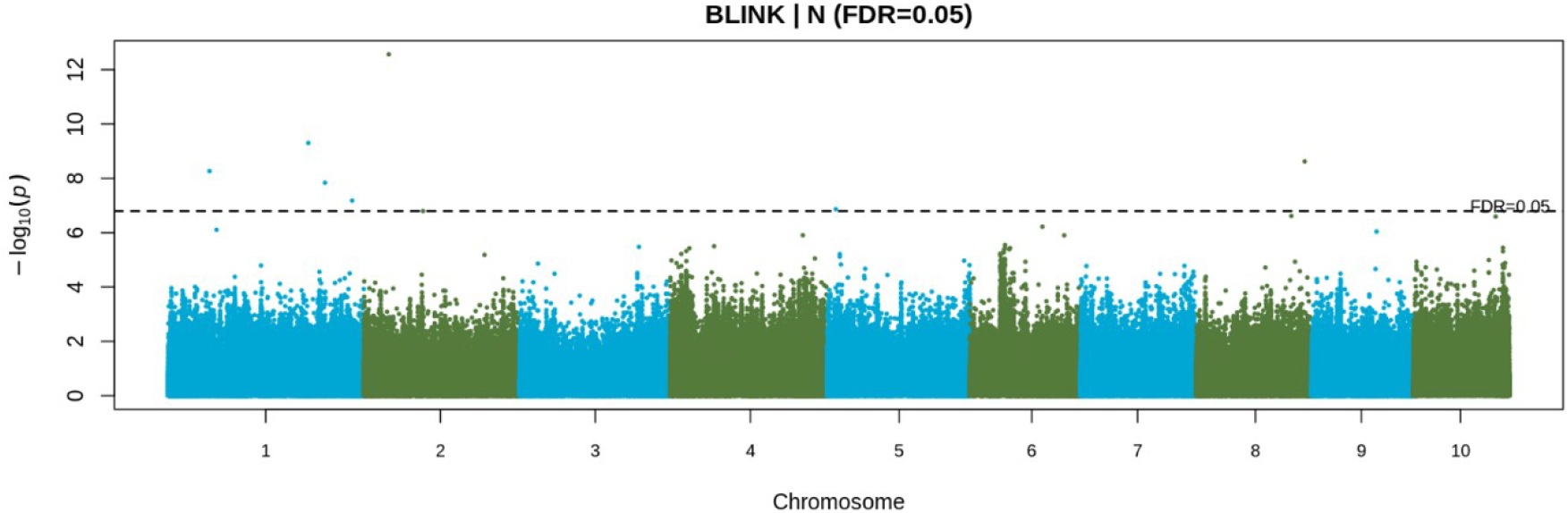
Manhattan plot of BLINK results for absolute free Asn. Points above dashed line represents results that passed false discovery rate correction of 5%.

**Fig. S7.**
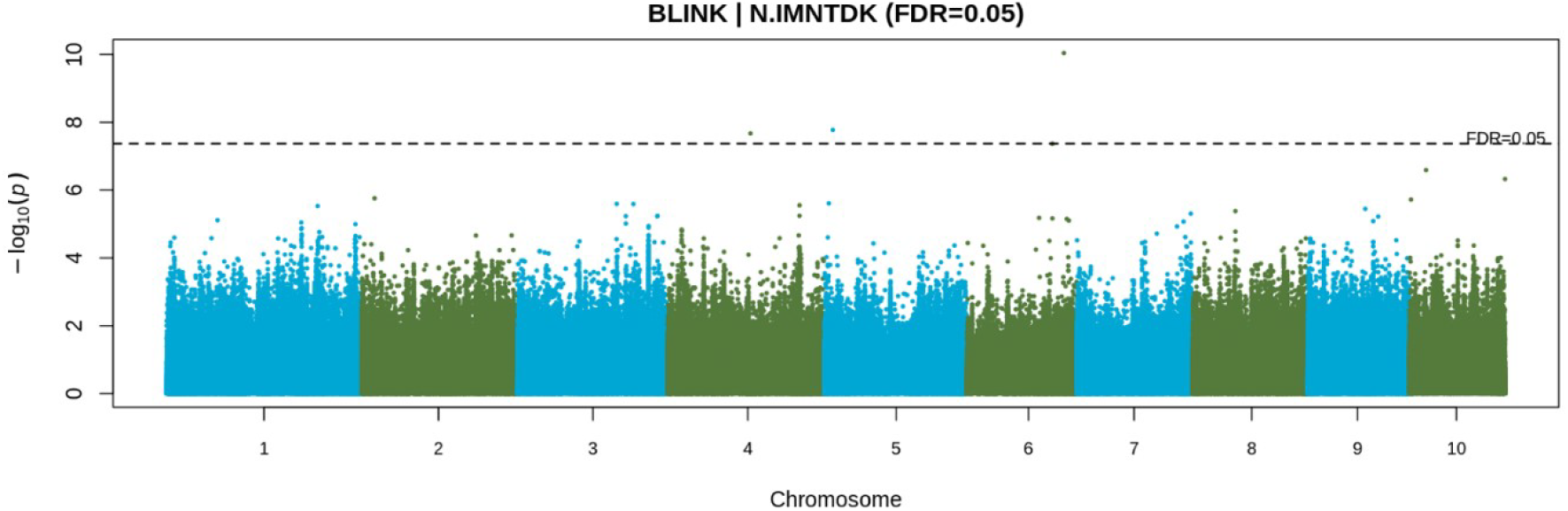
Manhattan plot of BLINK results for free Asn relative to aspartate family amino acids. Points above dashed line represents results that passed false discovery rate correction of 5%.

**Fig. S8.**
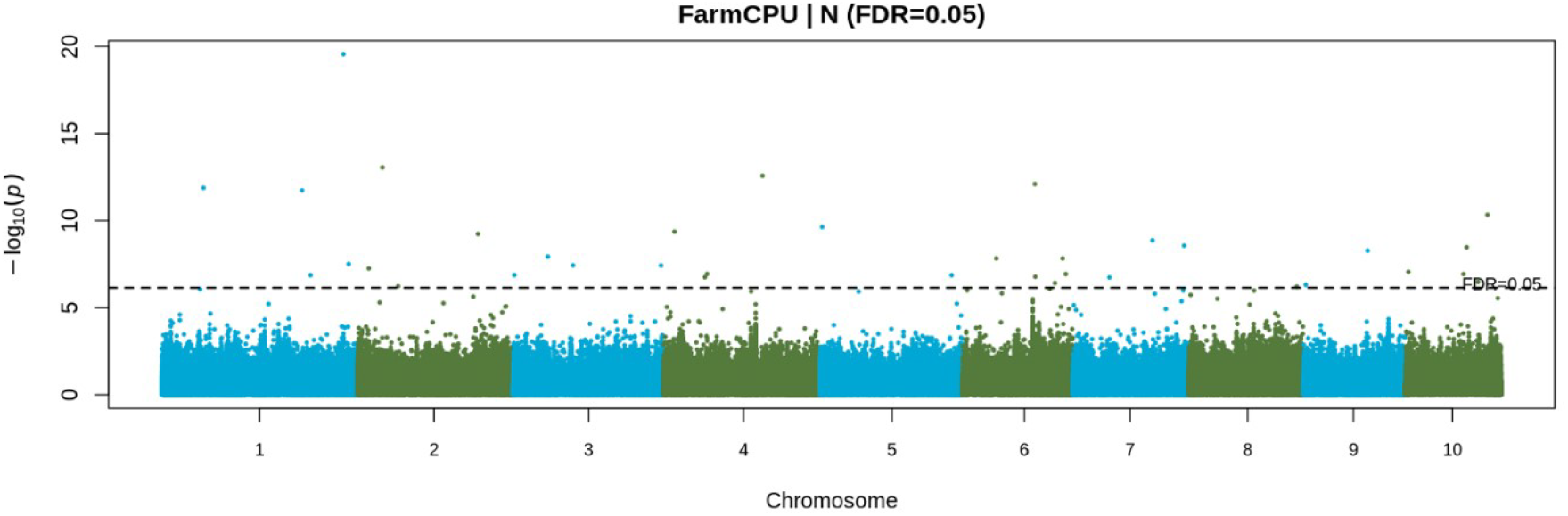
Manhattan plot of FarmCPU results for absolute free Asn. Points above dashed line represents results that passed false discovery rate correction of 5%.

**Fig. S9.**
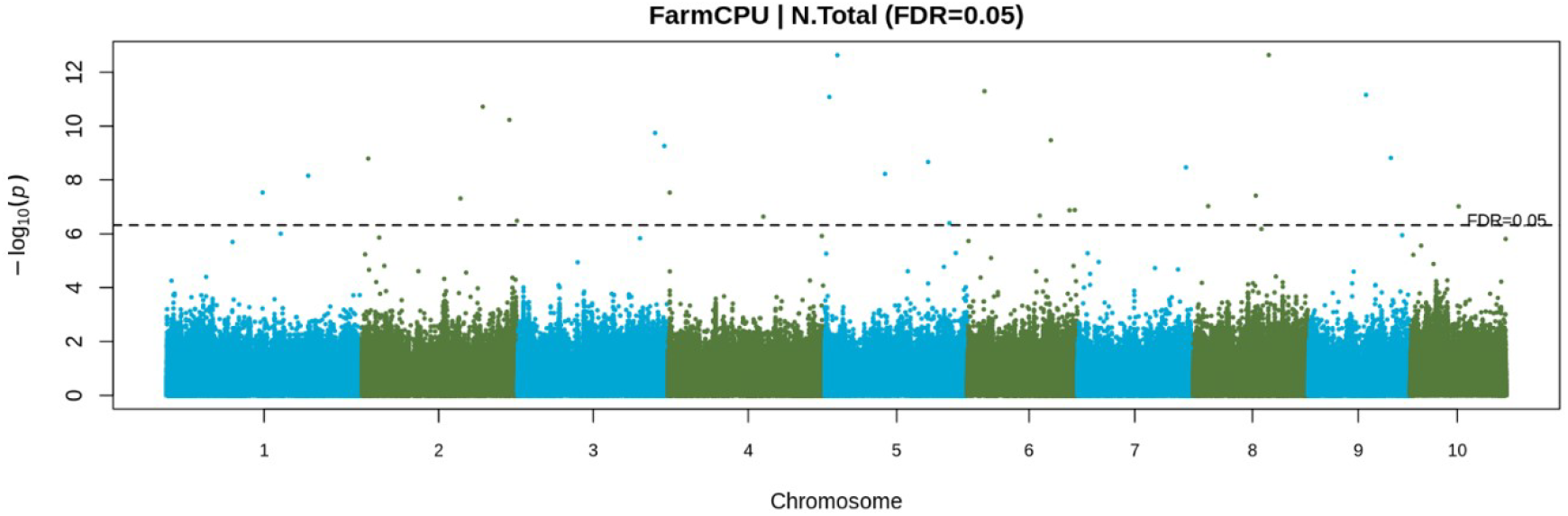
Manhattan plot of FarmCPU results for free Asn relative to total free amino acids. Points above dashed line represents results that passed false discovery rate correction of 5%.

**Fig. S10.**
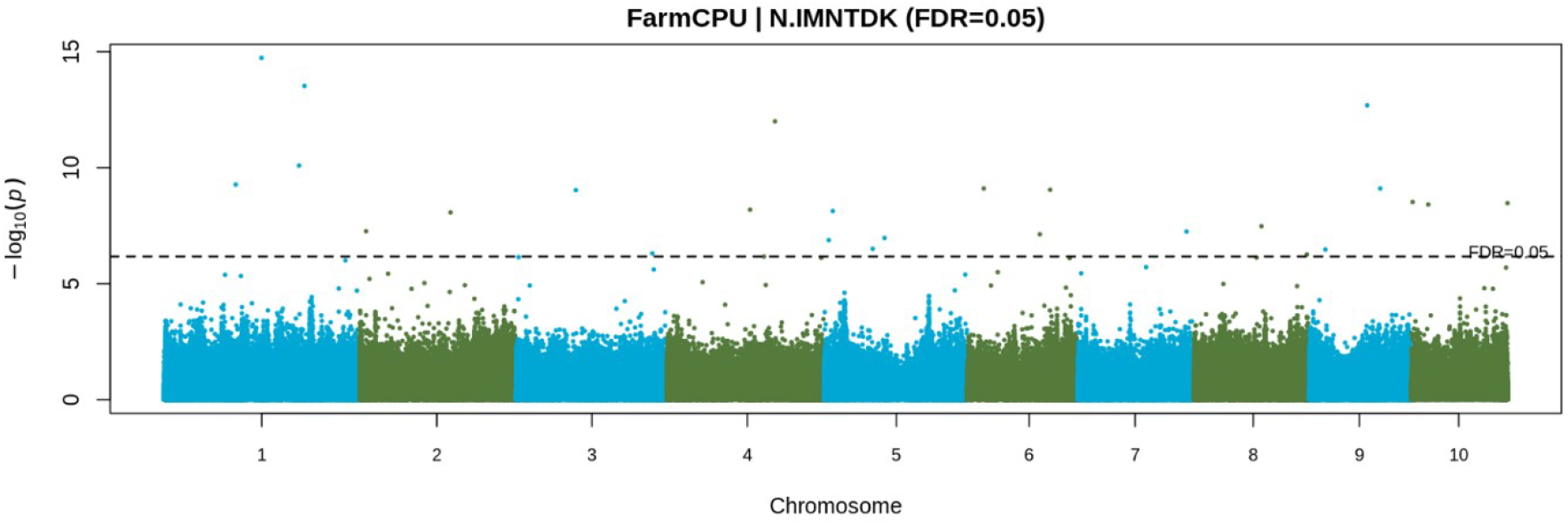
Manhattan plot of FarmCPU results for free Asn relative to aspartate family free amino acids. Points above dashed line represents results that passed false discovery rate correction of 5%.

**Fig. S11.**
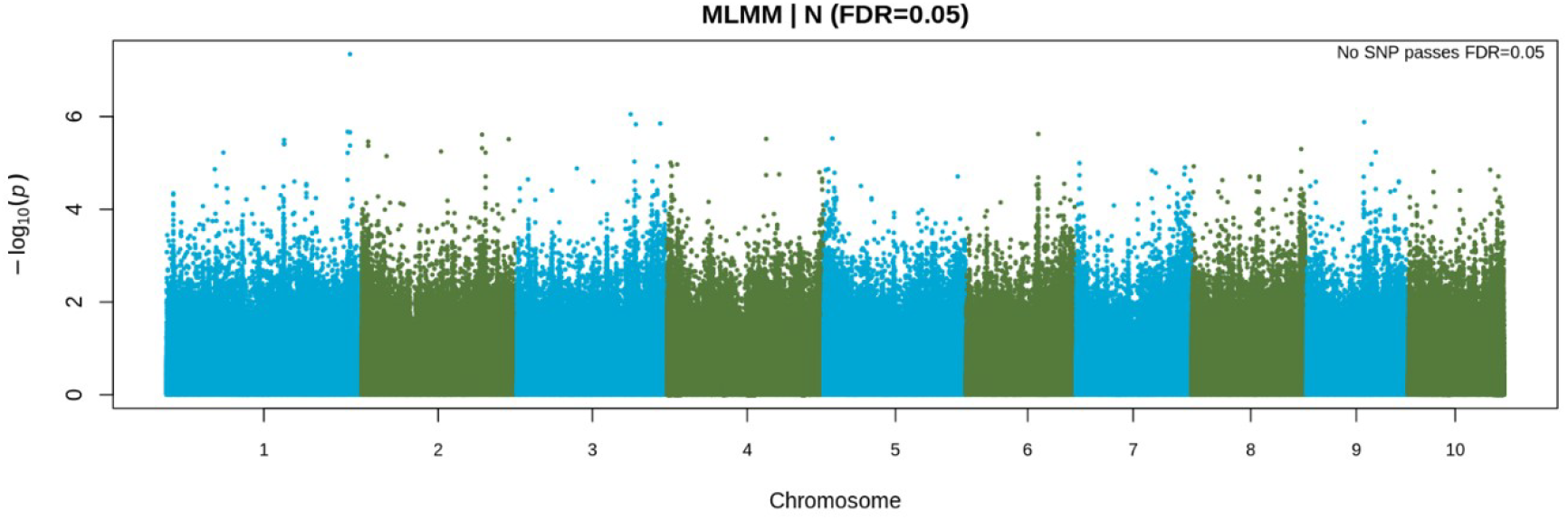
Manhattan plot of MLMM results for absolute free Asn. No SNP passes false discovery rate correction of 5%.

**Fig. S12.**
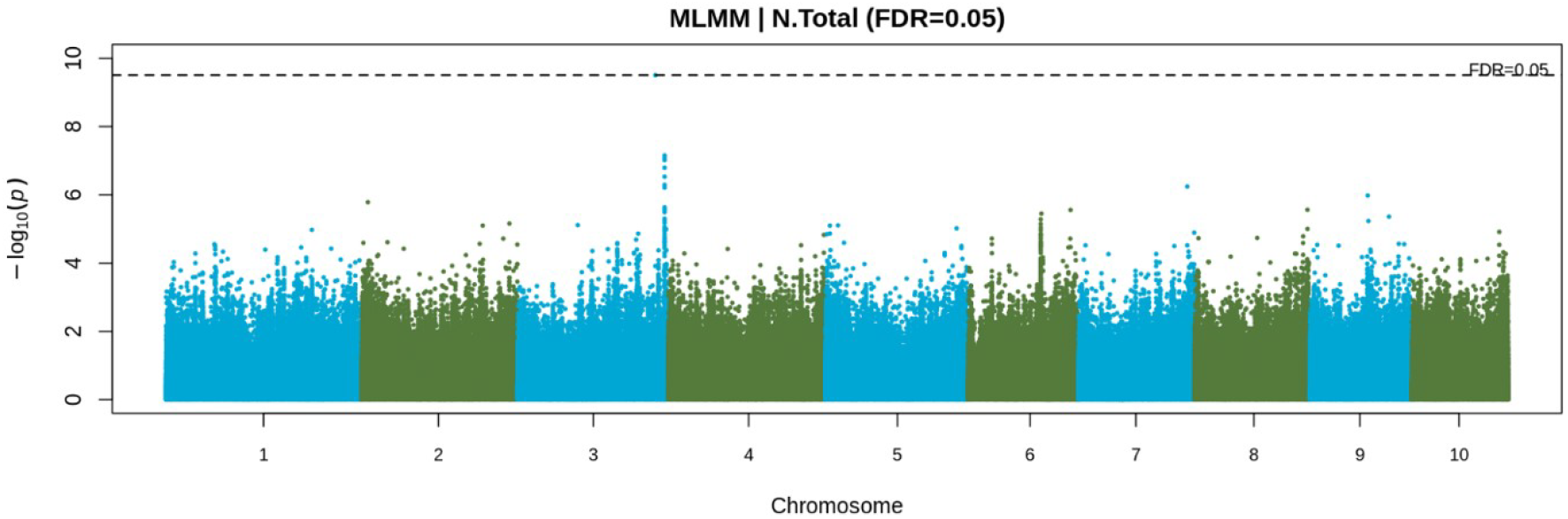
Manhattan plot of MLMM results for free Asn relative to total free amino acids. Points above dashed line represents results that passed false discovery rate correction of 5%.

**Fig. S13.**
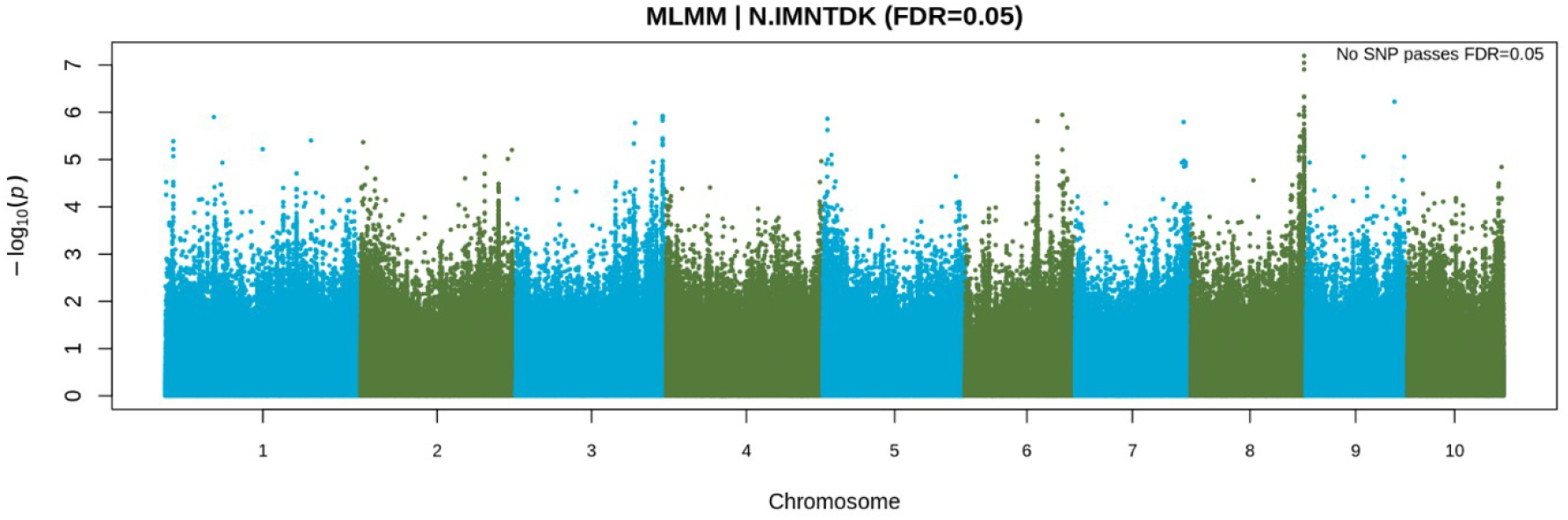
Manhattan plot of MLMM results for free Asn relative to aspartate family amino acids. No SNP passes false discovery rate correction of 5%.

